# Assessing the diversity and functional profile of the “microbial proteome” in fermented foods

**DOI:** 10.1101/2025.11.19.689333

**Authors:** Laura Winkler, Ayesha Awan, Nicole M. Rideout, Manuel Kleiner

**Affiliations:** Department of Plant and Microbial Biology, North Carolina State University, Raleigh, NC

## Abstract

Fermented foods are staples in diets worldwide and are known for their health benefits. Microorganisms are the key to fermented food production as they convert raw substrates into digestible, nutritious, and health-promoting products. While microbes are essential for fermented food production, their contribution to the dietary protein profile of the final food product in terms of microbial biomass is largely unknown. We analyzed proteins from 17 fermented foods using metaproteomics to identify and quantify microbial and food-derived proteins. We found that microbial proteins contribute up to 11% of the total protein content in fermented foods and comprise as much as 60% of the total number of identified proteins. These microbial proteins included many for central functions in microbial cells, such as glycolysis enzymes, translation machinery, and chaperones, as well as proteins for specialized functions that are important for the ecological niches in food fermentation, such as carbohydrate degrading enzymes and proteases. Some of these microbial proteins, such as proteases, could impact gut physiology. These findings highlight the substantial contribution of microbial proteins to the nutritional and functional profile of fermented foods, which may have important implications for interactions with the gut microbiota and health outcomes.

## Introduction

Fermented foods have been an integral part of the human diet for more than 10,000 years. They are consumed globally due to their enhanced shelf life, improved sensory properties, and health benefits. They include foods and beverages that are made through microbial conversion of the raw food substrate.^1–4^ While originally fermentation processes were mainly used as a food preservation method and to improve the taste and texture of certain foods, fermented products account for up to 40% of global food consumption today.^5,6^ This includes different kinds of bread, meat, cheese, alcoholic beverages, yogurt, sauerkraut, kimchi, and miso. For instance, the average annual consumption of plain bread in Turkey is 104 kg/person, miso in Japan is 7 kg/person, and yogurt in the Netherlands is 25 L/person.^5,7^

Beyond preservation and taste enhancement, fermentation improves the nutritional and functional profile of foods. It can enhance protein digestibility and micronutrient bioavailability in plant-based foods, particularly legumes, cereals, and pseudocereals.^8^ Proteins can also be hydrolyzed into bioactive peptides with antioxidant, antimicrobial, anti-diabetic, and anti-cancer properties.^9,10^ Fermented foods are also recognized as sources of probiotics and bioactive compounds, such as lactic acid and B vitamins, with potential benefits for gut health and immune modulation.^1,4,11^ While clinical studies link the consumption of fermented foods like yogurt and kimchi to reduced risks of cardiovascular disease, type 2 diabetes, and overall mortality, evidence supporting the effectiveness of most fermented foods for gastrointestinal health and disease is currently quite limited.^12–14^

One key aspect to consider when uncovering the mechanisms underlying the beneficial health effects of fermented foods is the microbial biomass, and in particular microbial protein, consumed as part of the fermented foods. When consuming fermented foods, it is often assumed that the dietary protein is mainly from the raw substrate and microbial proteins are not given much consideration. The extent to which proteins in the raw food substrate are converted into microbial proteins, and the identities of those microbial proteins, are unknown. Microbial biomass generated during fermentation may significantly contribute to the total protein content ingested during fermented food consumption. The contribution of microbes to fermented foods is rarely quantified or characterized, and their nutritional and functional roles are underexplored.

While various -omics approaches have been applied to study the microbiome of fermented foods, metaproteomics, the large-scale study of proteins from microbial communities, remains underutilized.^15–17^ Liquid chromatography–tandem mass spectrometry (LC-MS/MS) has only recently gained traction in food proteomics^18^, highlighting a knowledge gap about the protein composition of fermented foods, particularly the role of microbial proteins.

To determine the contribution of microbial proteins to fermented foods and estimate how much of the input substrate is converted to microbial biomass, we performed metaproteomic analyses on 17 fermented and 3 non-fermented food sources, including dairy products, sourdough bread, fermented cabbage, and cacao. We found that microbial proteins comprised a large portion of the total number of identifiable proteins in fermented foods. These microbial proteins make a substantial contribution to the total dietary protein content of these fermented foods and may possess distinct functional properties relevant to host health.

## Methods

### Sample acquisition

We selected suitable fermented foods based on products commonly mentioned in the literature.^19,20^ The foods were purchased from different suppliers or donated by people who make specific fermented foods in their homes. We also purchased three different non-fermented food sources, which serve as negative controls (NC). The selected foods are described in Table 1 and Supplemental Table 1.

**Table 1.** Fermented food selection and brand. The left column shows the selected fermented food and the right column indicates the brand (if purchased) or Homemade. NC: Negative, non-fermented control

| Food source | Brand or Homemade |
| --- | --- |
| Dairy yogurt | Siggi's |
| Dairy sour cream | Daisy |
| Cottage cheese | Good culture |
| Plant-based yogurt | So delicious |
| Plant based sour cream | Forager project |
| Miso paste | Miso Master<sup>®</sup> |
| Tempeh | Lightlife |
| Plain yeast bread | The essential baking company |
| Sourdough bread purchased | EuroClassic |
| Sourdough starter | Homemade |
| Sourdough bread | Homemade |
| Fermented Cabbage | Homemade |
| French Brie Cheese | Boar's head |
| Soy sauce | Kikkoman |
| Cacao | NavitasOrganics |
| Kefir day 1 fermented | Homemade |
| Kefir day 2 fermented | Homemade |
| Tofu (NC) | House Foods |
| Wheat flour (NC) | White Lilly |
| Cow milk (NC) | Simple Truth Organic |

### DNA extraction

We extracted DNA from each food sample using the DNeasy^Ⓡ^ PowerSoil^Ⓡ^ Pro kit (Qiagen GmbH, Germany) following the manufacturer’s instructions with small changes. 250 mg of each sample was loaded into the provided PowerBead tubes and 800 μL of CD1 solution was added. Cell lysis and homogenization was done using the BEAD RUPTOR ELITE (OMNI INTERNATIONAL) at 6 m/s, 00:45 min run, 00:30 min dwell, 3 cycles. For cow milk and tofu, this step was repeated 2 times, and for cacao, fermented cabbage, and wheat flour, this step was repeated 3 times. The samples were centrifuged at 15,000 x g for 1 min, and 200 µL of solution CD2 was added to the supernatant and mixed. Samples were centrifuged again and 600 µL of solution CD3 was mixed with the supernatant. 650 µL of lysate was added to a MB spin column, the column was centrifuged at 15,000 x g, and the flow-through was discarded. This step was repeated until all the lysate had been applied to the column. The column was placed into a new collection tube and 500 µL of solution EA was added, followed by centrifugation. The flow-through was discarded, 500 µL of C5 solution was added, and the column was centrifuged again. After that, the column was placed into a new collection tube and centrifuged for 2 min. Then, the column was placed into a 1.5 mL elution tube, and 100 µL of solution C6 was added to the center of the filter membrane. After centrifugation for 1 min, the eluted DNA was ready for downstream processing. The DNA quantity and quality were measured using the DeNovix^Ⓡ^ DS-11 FX + Spectrophotometer/Fluorometer (DeNovix Inc.). Extracted DNA was stored at −20°C.

### Library Preparation and Sequencing Methods

Sequencing was done by CosmosID (CosmosID Inc.). DNA libraries were prepared using the Watchmaker DNA Library Prep Kit (7K0019-1K) and compatible Twist Universal Adapter System. Genomic DNA was fragmented using a mastermix of Watchmaker Frag/AT Buffer and Frag/AT Enzyme Mix. Twist Universal Adapters (10X) and Twist HT Unique Dual Indexes (2X) were added to each sample, followed by 7 cycles of PCR to construct the DNA libraries. The final DNA libraries were purified using AMPure magnetic beads (Beckman Coulter) and eluted in nuclease-free water. Following elution, the libraries were quantified using the Qubit™ fluorometer dsDNA HS Assay Kit. Libraries were then circularized using the Element Adept library compatibility workflow and sequenced on the Element AVITI platform using the AVITI 2x150 Cloudbreak sequencing kit.

### Post sequencing data analysis

We analyzed the quality of raw reads using FastQC^21^, and merged forward and reverse reads using the reformat.sh function in BBTools.^22^ We trimmed adapters and removed contaminant phiX174 sequences using Trim Galore. We used phyloflash to map reads to 16S rRNA gene sequences and determine which taxa were present in the samples.^23^

### Sample preparation and protein extraction

We lyophilized all liquid food samples, including cow milk, soy sauce, and kefir, to concentrate them. We weighed 150 mg of wet sample (such as yogurt) and 100 mg of dry sample (such as bread) into 2 mL Lysing Matrix E (MP Biomedicals™) tubes and added 1 mL of SDS-lysis buffer (5% w/v) in 50 mM TEAB, pH 7.5. We lysed samples using a BEAD RUPTOR ELITE (OMNI INTERNATIONAL) at the following settings: 6.45 m/s, 00:45 min run, 01:00 min dwell, 5 cycles. Subsequently, the samples were heated for 10 min at 95°C and then centrifuged for 5 min at 21,000 x g and the supernatant was retained.

To extract proteins and prepare peptides, we followed the Suspension Trapping (S-Trap) sample preparation method for bottom-up proteomics analysis.^24^ We mixed 6.4 µL of 500 mM DTT with the entire lysate and incubated for 10 min at 95°C. Then, we added 12.7 µL of 500mM IAA solution and incubated in a centrifuge tube shaker at room temperature (RT) and 600 rpm for 1 min. The lysates were then further incubated in the dark for 30 min. After the incubation, 16 µL of 12%-phosphoric acid solution and 6 times the sample volume of binding/wash buffer (100mM TEAB in 90% MetOH) was added.

For the next steps, the flow-through was discarded after every centrifugation step. 1200 µL of sample was loaded on the S-TRAP-mini-80 column (PROTIFI^TM^) and centrifuged at 4,000 x g for 30 sec. The column was washed with 400 µL of wash/binding buffer and centrifuged at 4,000 x g for 30 sec. This step was repeated 3 times in total. Without adding more buffer, the column was centrifuged one more time at 4,000 x g for 30 sec. The column was then placed in a new 2 mL sample tube, and proteins were digested by adding 0.8 µg MS-grade trypsin (Thermo Scientific Pierce^TM^, Rockford, IL, USA), solubilized in 40 µL digestion/elution buffer 1 (50 mM TEAB in HPLC water) to the column and incubating it overnight at 37°C in a wet chamber. The next day, peptides were eluted off the column by first adding 80 µL of elution buffer 1 to the column, incubating for 10 min at 37°C, and centrifuging at 4,000 x g for 1 min. Then 80 µL of elution buffer 2 (0.2% formic acid in HPLC water) was added and centrifuged at 4,000 x g for 1 min. Finally, 80 µL of elution buffer 3 (50% acetonitrile and 0.2% formic acid in HPLC water) was added and centrifuged at 4,000 x g for 1 min. Acetonitrile from eluted peptides was removed using a vacuum centrifuge and the peptides were resuspended in ∼ 30 µL elution buffer 2. Peptide concentrations were determined using the Micro BCA^TM^ Protein Assay Kit (Thermo Scientific) according to the manufacturer’s instructions.

### LC-MS/MS analysis

We analyzed peptides from food samples using 1D-LC-MS/MS as previously described.^25,26^ We loaded 1 μg of peptides onto a 5 mm, 300 μm ID C18 556 Acclaim® PepMap100 pre-column (Thermo Fisher Scientific) with loading solvent A (2% acetonitrile, 0.05% TFA) using an UltiMate™ 3000 RSLCnano Liquid Chromatograph (Thermo 558 Fisher Scientific). An EASY-Spray analytical column heated to 60°C (PepMap RSLC C18, 2 μm material, 75 cm × 75 μm, Thermo Fisher Scientific) was used to separate the peptides. A 140 min gradient at a flow rate of 300 nl/min was used for peptide separation of which the first 102 minutes of the gradient went from 95% eluent A (0.1% formic acid) to 31% eluent B (0.1% formic acid, 80% acetonitrile), then 18 min from 31 to 50% B, and 20 min at 99% B. One to two 100% acetonitrile wash runs were performed between each sample to minimize carryover. The eluted peptides were ionized using an Easy-Spray source and analyzed in an Exploris 480 hybrid quadrupole-Orbitrap mass spectrometer (Thermo Fisher Scientific) with the following parameters: m/z 445.12003 lock mass, normalized collision energy 24, 25 s dynamic exclusion, and exclusion of ions of +1 charge state. Full MS scans were acquired for 380 to 1600 m/z at a resolution of 60,000 and a maximum IT time of 200 ms. Data-dependent MS^2^ was performed for the 15 most abundant ions at a resolution of 15,00 and maximum IT of 100 ms.

### Database construction and protein identification

For each of the fermented foods we constructed non-redundant protein sequence databases following the previously described principles.^27^ These databases comprised annotated protein sequences from the Proteomes section of UniProt corresponding to the bacterial and eukaryotic ingredients present in each food source. The database for each food source is unique, and the species to include in the database were selected based on the current literature, the ingredients list of the food, and our taxonomic analysis of the metagenomic data. The database composition for each food source is detailed in Supplemental Table 2. In order to remove redundant sequences, the protein sequences of each strain/species were clustered using cd-hit (Version 4.7) with an identity threshold of 95%.^28^

We searched the MS^2^ spectra against the respective food-source-specific database using the PEAKS^®^ X + software and quantified proteins using area under the curve^29–31^ with the following settings: Precursor correction: Min charge=1, Max charge=10; Precursor mass error tolerance=10 ppm using monoisotopic mass; Fragment ion error tolerance=0.05 Da; Enzyme=trypsin, semispecific; maximum missed cleavages=2; Maximum allowed variable PTMs per peptide=3, including Carbamidomethylation, Oxidation (M) and Deamidation (NQ). Advanced options: DeNovo Sequencing, FDR decoy-fusion, Identification of unspecified PTMs with PEAKS PTM and Identification of more mutations with PEAKS SPIDER. For the downstream analysis protein tables were filtered for a FDR of 5%, Proteins -10lgP ≥ 15, and ≥ 2 unique peptides without significant peptides and a DeNovo only score of 50% was used.

### Data processing, statistical analysis, and data visualization

We used total sum scaling to (1) normalize the relative abundances of proteins within each sample^32^ and to (2) normalize at the organism level to estimate the relative abundances of proteins within each organism in a sample. We calculated the abundance of each microbial species using its proteinaceous biomass.^33^ We did this by summing up the peptide spectra of all proteins with at least 2 protein unique peptides assigned to each given species. GraphPad Prism (Version 10.2.3), Microsoft Excel and ggplot2^34^ in R (version 2024.12.1)^35^ were used for data processing and visualization. The pheatmap package in R was used to make the heatmap.^36^

## Results

### Microbial proteins comprise a substantial portion of the total protein content in fermented foods

We used metaproteomics to analyze the protein content of 17 different fermented foods and 3 non-fermented foods (Fig.1). The total number of identified proteins in each food source ranged from 680 total proteins in dairy sour cream to beyond 3,000 total proteins in homemade sourdough bread (Fig.1A). The number of identified microbial proteins ranged from 119 proteins in Miso to over 1,000 different proteins in Brie cheese. The relative abundance of all microbial proteins out of the total detected proteins ranged from 0.6% in cacao to more than 11% in plain yeast bread (Fig. 1B).

**Figure 1.**
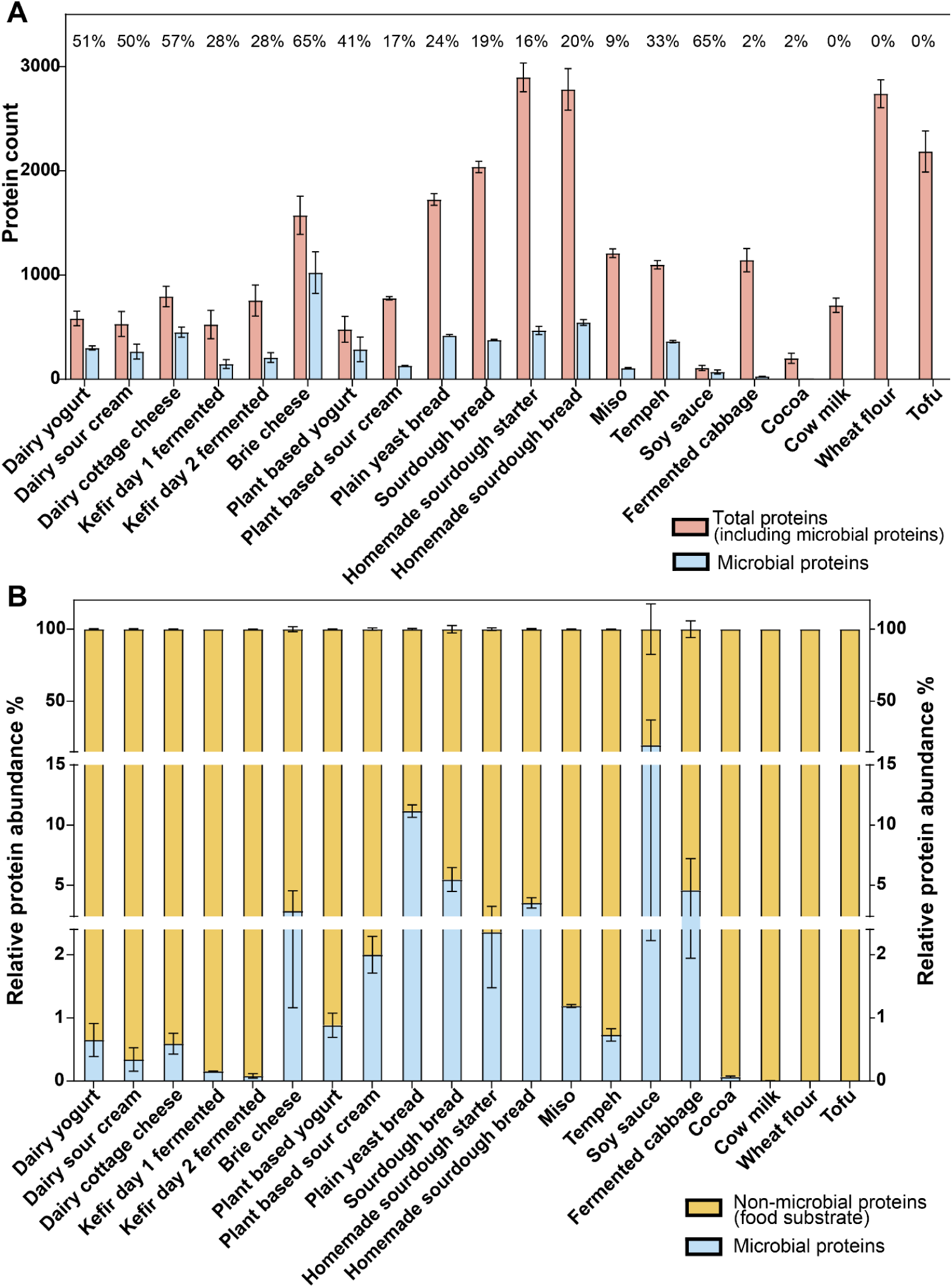
Microbial proteins contribute a large amount of dietary protein to fermented foods. **A.** Contribution of the different identified microbial proteins to the overall number of identified proteins (mean and standard deviation are shown, 3 technical replicates were analyzed). The percentage above each food source indicates the proportion of identified microbial proteins relative to the total number of proteins detected. **B.** Relative abundance of microbial and non-microbial (food substrate) protein in different fermented food sources (n=3). Cow milk, Wheat Flour and Tofu served as unfermented negative controls. Peptides were identified using the PEAKS software, allowing for semi-tryptic peptides. Relative proteinaceous biomass was determined as described in Kleiner et al., 2017 for each sample and the results were averaged by food.^33^

The proportion and diversity of microbial proteins was much higher than the food substrate proteins in 5 out of the 16 fermented foods analyzed. In Brie cheese, for example, out of the 1,573 different proteins present, 1,023 of these (65%) were microbial proteins. This pattern was observed in almost all dairy products, especially in dairy yogurt, dairy sour cream, and dairy cottage cheese, with the exception of kefir, in which microbial proteins only made up 23-27% of the total number of proteins. In plant-based dairy alternative fermented foods, the number of microbial proteins ranged from 16% of all proteins in plant-based sour cream to 43% in plant-based yogurt. In the bread samples, microbial proteins contributed 17-19% to the overall protein number. The proportion of microbial proteins by number was lower in miso (9%) and fermented cabbage (2%). In contrast, 32-57% of the overall number of proteins in tempeh and soy sauce were of microbial origin.

Looking at total protein biomass regardless of protein identity, the percentage of each sample that could be attributed to microbial protein biomass varied between different fermented foods with 0.5-2.5% in dairy products, 0.8-2% in plant-based dairy alternatives, 0.5-1.2% in soy based products, 5% in cabbage and 5-11% in bread products. These results show that microbial proteins contribute a large proportion of the protein, by both number of different proteins and abundance relative to non-microbial proteins, to the overall protein content of fermented foods.

### Fermented foods harbor distinct, food-specific microbial communities

As expected, microbial community composition differed strongly between dairy and plant-based fermented foods, as well as between different dairy products such as yogurt and cheese (Fig. 2). Among all the fermented foods analyzed, brie cheese and kefir had the highest number of microbial species, while plain yeast bread and soy sauce had the lowest. While both dairy and plant yogurts contained *Streptococcus thermophilus*, *Lacticaseibacillus paracasei, Lactobacillus delbrueckii* ssp*. bulgaricus* and *Lactobacillus acidophilus,* only dairy yogurt contained *Bifidobacterium animalis* and only plant yogurt contained *Lacticaseibacillus rhamnosus*. Dairy and plant sour creams had distinct microbial compositions and dairy sour cream clustered with dairy cottage cheese, while plant sour cream clustered with plant and dairy yogurts. Dairy sour cream contained *Lactococcus lactis* strains and *Leuconostoc mesenteroides,* while plant sour cream contained *L. acidophilus*, *L. delbrueckii* and *S. thermophilus*.

**Figure 2.**
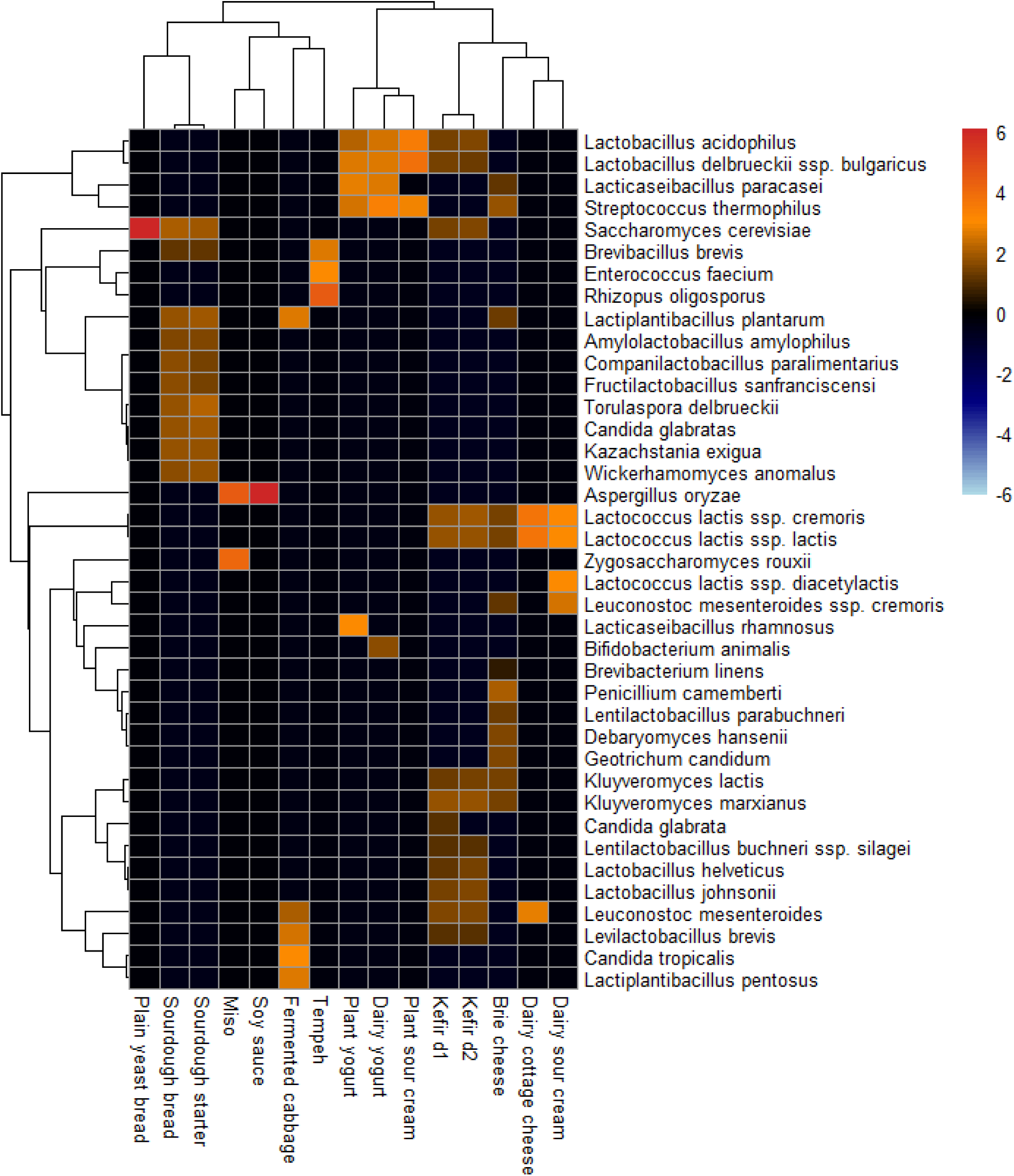
Microbial community composition varies by fermented food. Heatmap depicting the log2-transformed relative abundances of microbial taxa in different fermented foods measured using metaproteomics. The relative abundance for each strain was z-scored across all food samples, which shows how much more or less abundant that strain is in a given food compared to its average abundance across all foods. Three technical replicates were measured and averaged for each food type. The underlying data are available in Supplemental Table 3.

Although both dairy-based fermented foods and their plant alternatives mainly contained Bacillota species, the specific taxa differed. Brie cheese and kefir had the greatest diversity of microbes compared to other dairy fermented foods and contained both bacterial and fungal phyla. Bacterial taxa in brie cheese included the Bacillota species *L. lactis* ssp. *lactis, L. lactis* ssp. *cremoris,* and *S. thermophilus,* as well as Actinobacteria species *Brevibacterium linens*. Fungal taxa in brie cheese included Ascomycota species *Kluyveromyces lactis, Debaryomyces hansenii, Penicillium camemberti,* and *Geotrichum candidum*.

Kefir, a fermented milk product, contained the fungal species *Candida glabrata, Kluyveromyces marxianus, K. lactis* and *Saccharomyces cerevisiae* as well as the bacterial species *L. acidophilus, Lactobacillus johnsonii, Lactobacillus helveticus, Lentilactobacillus buchneri, L. mesenteroides, L. delbrueckii* and *L. lactis*. Kefir fermented for 2 days, compared to 1 day of fermentation, had a higher abundance of *L. johnsonii* and *L. helveticus* and a lower abundance of *C. glabrata*. Legume (soy) based fermented foods like tempeh, soy, and miso were dominated by fungal species, including *Rhizopus oligosporus* in tempeh, *Zygosaccharomyces rouxii* in miso, and *Aspergillus oryzae* in both miso and soy sauce. Tempeh also contained the bacterial species *Enterococcus faecium* and *Brevibacillus brevis*.

Fermented cereal-based foods, including plain yeast bread and sourdough bread, both contained *S. cerevisiae*. However, sourdough contained a greater diversity of both Ascomycota fungal species, including *Wickerhamomyces anomalus, C. glabrata, Torulaspora delbrueckii, Kazachstania exigua,* and Bacillota bacterial species, including *Lactiplantibacillus plantarum, Amylolactobacillus amylophilus, Companilactobacillus paralimentarius, Fructilactobacillus sanfranciscensis* and *B. brevis*. Fermented cabbage or sauerkraut, another plant-fermented food, also contained fungal and bacterial species, including *Candida tropicalis, Lactiplantibacillus pentosus, L. plantarum* and *Levilactobacillus brevis*.

The overall microbial community composition was substrate-specific, with both dairy and plant-fermented foods dominated by Bacillota and Ascomycota species. *L. delbrueckii* ssp*. bulgaricus, L. lactis* and *L. acidophilus* were specific to mainly dairy fermented products and *Lactiplantibacillus* species were associated with plant substrates containing complex carbohydrates. Fungal species were mainly associated with legume and cereal-based fermented foods.

### Distinct profiles of microbial and non-microbial proteins across various fermented foods

We quantified the relative abundances of microbial and food substrate-derived non-microbial proteins across 14 dairy, plant, grain, and legume based fermented foods, as well as their unfermented food substrates (Figure 3). In most of the fermented food samples, the diversity of microbial proteins was greater than that of non-microbial, food substrate dietary proteins. Comparison with the unfermented food counterparts revealed that microbial fermentation significantly alters the protein composition of these foods.

**Figure 3.**
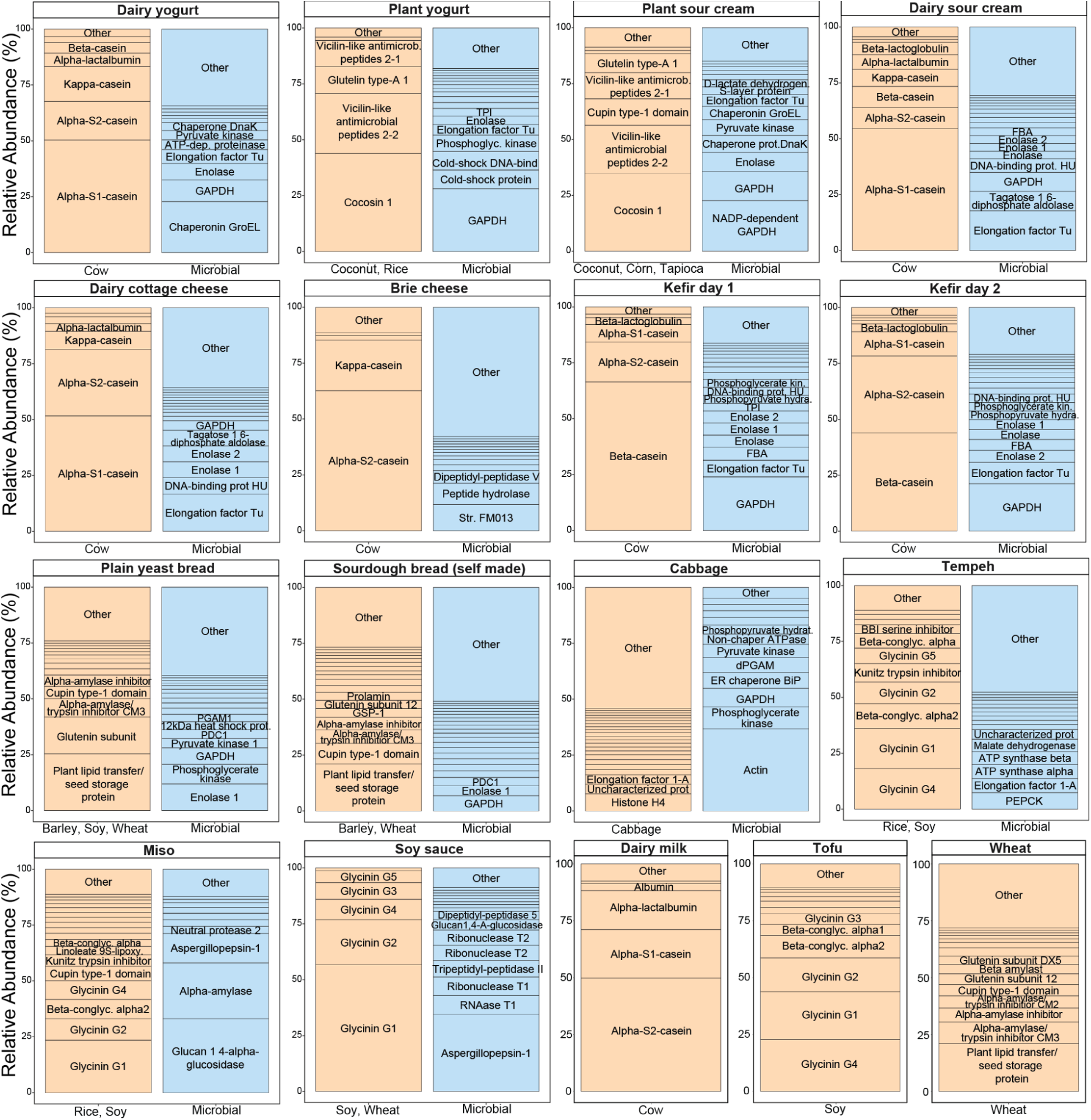
Distribution of microbial and non-microbial proteins in fermented foods. Stacked bar plots showing relative abundances and diversity of the microbial and food substrate-derived non-microbial proteins in different fermented foods. Proteins that were at least 1% of the proteome were plotted in distinct stacks, while low abundance proteins were summed in the “Other” category. Proteins with at least 3% relative abundance are labelled. Abundances are an average across three replicates and microbial and non-microbial protein abundances were normalized separately to 100% within each category. The underlying data are available in Supplemental Data Table 4. Abbreviated protein names are as follows: GAPDH (glyceraldehyde-3-phosphate dehydrogenase), PEPCK (phosphoenolpyruvate carboxykinase), PDC1 (pyruvate decarboxylase 1) and FBA (fructose-1,6-bisphosphate aldolase).

In unfermented dairy milk alpha-S2-casein, alpha-S1-casein and alpha-lactalbumin were the most abundant non-microbial proteins. In fermented dairy products such as yogurt and cottage cheese, alpha-S1-casein, alpha-S2-casein, kappa casein and alpha lactalbumin were the most abundant cow-derived proteins, while beta-casein and beta lactalbumin were higher in abundance in kefir and sour cream. Brie cheese had a high abundance of alpha-S2-casein and kappa casein.

Microbial proteins in dairy fermented foods varied by product type. Yogurt had a high abundance of chaperone proteins, glyceraldehyde-3-phosphate dehydrogenase (GAPDH) and enolase. Cottage cheese and sour cream had a high abundance of elongation factor Tu, DNA-binding protein HU, enolases and tagatose-1,6-diphosphate aldolase. Kefir had a high abundance of GAPDH, elongation factor Tu and enolase. Brie cheese had a high abundance of mainly fungal proteins, including peptide hydrolase and dipeptidyl-peptidase V.

Plant-based alternatives to dairy fermented foods had distinct profiles of non-microbial food substrate proteins compared to their dairy counterparts, including high abundance of cocosin 1, vicillin-like antimicrobial peptides, glutelin type A1 and cupin domain containing protein. However, the microbial proteins, including GAPDH, enolase, DNA-binding proteins, chaperones, and elongation factor Tu were the most abundant microbial proteins in plant-based and dairy yogurts and sour creams.

Unfermented wheat had a high abundance of seed storage proteins ( *e.g.,* 7.14% abundance of plant lipid transfer/seed storage protein), glutenin (10% summed abundance of all subunits), and a variety of amylase and trypsin inhibitors (>20% summed relative abundance), whereas its fermented counterparts, such as yeast bread and sourdough, maintained a high abundance of seed storage proteins (14% abundance of plant lipid transfer/seed storage protein) but had a lower abundance of amylase and trypsin inhibitors (∼14% summed relative abundance). Grain-based fermented foods, plain yeast bread, and sourdough had different microbial protein profiles reflective of differences in their microbial community and diversity (Figure 2). While both breads had abundant yeast-derived proteins with high abundance of GAPDH and enolase, sourdough had a much higher diversity of low-abundance bacterial proteins.

Similarly, unfermented soy-based tofu had a high abundance of seed storage proteins, including glycinin and beta-conglycinin, while fermented soy based products, including tempeh and miso, had reduced abundance of these seed storage proteins and increased abundances of protease inhibitors, including the kunitz trypsin inhibitor and the BBI serine inhibitor (Supplemental Data Table 4). In terms of microbial proteins, soy-based fermented food products had a high abundance of microbial carbohydrate degradation enzymes and proteases. Tempeh had a high abundance of microbial phosphoenolpyruvate carboxykinase (PEPCK), elongation factor 1-A, ATP synthase, and malate dehydrogenase. Miso had a high abundance of microbial glucan-1,4-alpha-glucosidase, alpha amylase, aspergillopepsin-1, and neutral protease 2.

Overall, these results indicate significant shifts in protein composition between fermented foods and their unfermented counterparts. Unfermented food substrate proteins, such as caseins, glutelins, vicilins, and glycinin, are reduced in relative abundance as they are degraded by microbes and used to produce new microbial proteins.

## Discussion

While it is known that fermented foods positively impact health outcomes, the underlying factors driving these effects are not well characterized.^37–39^ Thus, it is crucial to characterize the distinct microbial and non-microbial components of fermented foods to better understand how they might interact with the host and the gut microbiota upon consumption. While many different fermented foods and beverages from all over the world have been investigated using different -omics approaches, including metagenomics, metatranscriptomics and metabolomics, the metaproteomics approach remains underutilized despite the fact that proteins are key nutritional and functional components of these foods.^40–42^

In this study, we used metaproteomics to study fermented foods on the protein level, infer specific microbial and non-microbial proteins and characterize their functional profiles. Our metaproteomics results show that microbial proteins comprise a substantial proportion of total protein content in fermented foods. This shows that microorganisms not only contribute to the fermentation process itself but also to the overall nutritional profile of fermented food through the conversion of substrate protein into microbial protein. These results offer avenues for investigation beyond the known probiotic effects of fermented foods, as the microbial proteins consumed as part of these foods may have direct nutritional and immune-modulatory effects. The large microbial protein diversity present in these foods also translates to a broader functional potential, including carbohydrate-degrading enzymes, proteases, bioactive peptides and other compounds that can influence the host’s digestion and interact with the gut microbiota in the colon.^43,44^

As expected, the most abundant microbial proteins found in fermented foods were associated with central metabolism and core physiological functions; these proteins included glycolysis proteins, translation factors, chaperones, carbohydrate degrading enzymes and proteases.^40,41^ In foods fermented by bacteria, such as dairy foods, proteins involved in protein biosynthesis, carbohydrate metabolism and energy production were most abundant. In contrast, in foods that were mostly fermented by fungi, such as miso, soy sauce, and tempeh, carbohydrate-degrading enzymes (CAZymes) and proteases were more abundant .^45^

The presence of enzymes such as CAZymes and proteases is noteworthy as it points to substrate-specific adaptations that potentially impact gut health. CAZymes are crucial for the breakdown of carbohydrates and can support fiber breakdown, influencing gut community dynamics through cross-feeding.^46–48^ This indicates that fungi, such as *Aspergillus oryzae* (soy sauce and miso) and *R. oligosporus* (tempeh), potentially play a crucial role in carbohydrate accessibility to other microbes. Microbial proteases, found in brie cheese, soy sauce, miso, and tempeh, can act on substrates such as soy proteins to generate bioactive peptides that can affect health, including impacting angiotensin-converting enzyme (ACE) activity, nutrient processing and absorption, and modulating immune and gut barrier responses.^49–51^ While our results did not focus on profiling the functional activity of these health-relevant microbial proteins, future studies should investigate their activities and how they might interact with the host and other gut microbes.

In addition to increasing microbial protein biomass in fermented foods, microbial fermentation also altered the nutritional landscape of foods by modifying the abundance of functional and health-relevant food substrate proteins through metabolic conversion. These changes include degrading and altering the abundances of anti-nutritional protease inhibitors, such as alpha-amylase/trypsin inhibitors in wheat, Kunitz trypsin and BBI inhibitors in soy, as well as of potential food allergens and immunogenic proteins, including glutelin, vicillin and casein proteins.^8,52,53^ These findings have important implications in the context of previous studies showing that the detection and *in vitro* IgE reactivity of known food allergens is altered post microbial fermentation and highlight the potential for engineering the fermentation process to enhance microbial proteolytic activity against anti-nutritional factors or allergenic food proteins.^52–54^

Furthermore, the observation that different food types are dominated by different microbial taxa suggests that fermented foods carry a substrate-specific core functional microbiome. The identified taxa in the dairy products and plant-based products in this study aligned with findings of previous studies that used DNA-based approaches to profile microbes. The sourdough microbiome has been extensively studied, revealing that sourdough contains a large diversity of microorganisms, especially yeasts.^55–57^ We included the strains identified in those previous studies in the protein database for this study and our results based on proteinaceous biomass differed slightly. We found that *S. cerevisiae* was the dominant fungus, followed by *C. glabrata*, *K. exigua*, *T. delbrueckii* and *W. anomalus,* in line with previous studies.^58–60^ However, the most abundant lactobacillus bacteria in our sourdough were *L. plantarum* (9.7%), followed by *F. sanfranciscensis* (1.7%), while other studies have reported the reverse trend in the abundance patterns of these two bacteria using a 16S rRNA gene and ITS amplicon sequencing approach.^55^ This difference between our results and prior studies is likely due to the fact that metaproteomics measures the proteinaceous mass contribution of a species to a community, while DNA sequencing-based approaches provide insights into gene copy numbers contributed by species. This suggests that using proteinaceous biomass instead of DNA-based measures may be more representative of the quantitative profile of the microbial community in fermented foods, as well as its contribution to the nutritional and functional profile of the food.^33^

Overall, this study underscores the importance of considering microbial proteins as a key component of the nutritional and functional value of fermented foods. While the direct effects of some fungal and bacterial proteins and enzymes on human health remain speculative due to digestive degradation, their influence on the immunogenicity or functionality of other proteins warrants further investigation beyond the scope of this study. Our use of metaproteomics has laid the groundwork for such future research into the potential health-promoting effects of microbial proteins in fermented food products.

## Data availability

The mass spectrometry proteomics data have been deposited to the ProteomeXchange Consortium via the PRIDE partner repository with the dataset identifier PXD070912 [reviewer username; reviewer password: bwdXYjWqNeTb].^61^ The metagenomic raw reads were submitted to NCBI SRA under the bioproject identifier PRJNA1365831.

## Supplementary materials

**Supplemental Table 1:** Table containing details of each of the commercially obtained fermented foods, including source, label specified food substrate and microbes.

**Supplemental Table 2:** Table listing each of the microbial and non-microbial species used to construct the protein database that was used in the proteomics analysis.

**Supplemental Table 3:** Table containing the relative abundance of the proteomic biomass of each microbial species in the fermented foods analyzed.

**Supplemental Table 4:** Table containing the raw protein abundances identified in each of the three replicates of the fermented foods analyzed in the proteomic analysis.

## Author contributions

**Laura Winkler:** Experimental design, data analysis, writing and editing the manuscript

**Ayesha Awan:** Conceptualization of the study, experimental design, data collection, data analysis, writing and editing the manuscript

**Nicole Rideout:** Experimental design, data collection, data analysis, editing

**Manuel Kleiner:** Funding, conceptualization of the study, experimental design, data processing, writing and editing the manuscript

## Acknowledgements

We would like to thank Jessie Maier for providing the homemade sourdough starter and bread, Sadije Jakupi for providing the fermented cabbage and Sandra Kelberlau for providing the kefir. We would also like to thank Dr. Heather Maughan for editing the manuscript and for her insightful suggestions. All LC-MS/MS measurements were made in the Molecular Education, Technology, and Research Innovation Center (METRIC) at North Carolina State University. This work was supported by the National Institutes of Health under Awards R35GM138362 and R01DK118024, and the European Union under Award Number 2023 1 DE01 KA131 HED 000120145.

## References

1. Marco ML, Heeney D, Binda S, Cifelli CJ, Cotter PD, Foligné B, et al. Health benefits of fermented foods: microbiota and beyond. Current Opinion in Biotechnology. 2017 Apr;44:94–102.

2. Sahm H, Antranikian G, Stahmann KP, Takors R. Industrielle Mikrobiologie. 1st ed. Berlin, Heidelberg: Springer Berlin Heidelberg Imprint Springer Spektrum; 2013.

3. Tamang JP, Cotter PD, Endo A, Han NS, Kort R, Liu SQ, et al. Fermented foods in a global age: East meets West. Comp Rev Food Sci Food Safe. 2020 Jan;19(1):184–217.

4. Wilson DB, Sahm H, Stahmann KP, Koffas M, editors. Industrial microbiology. Weinheim: Wiley-VCH; 2020. 397 p.

5. Cuamatzin-García L, Rodríguez-Rugarcía P, El-Kassis EG, Galicia G, Meza-Jiménez MDL, Baños-Lara MaDR, et al. Traditional Fermented Foods and Beverages from around the World and Their Health Benefits. Microorganisms. 2022 June 2;10(6):1151.

6. Dimidi E, Cox S, Rossi M, Whelan K. Fermented Foods: Definitions and Characteristics, Impact on the Gut Microbiota and Effects on Gastrointestinal Health and Disease Nutrients. 2019 Aug 5;11(8):1806.

7. Chilton S, Burton J, Reid G. Inclusion of Fermented Foods in Food Guides around the World. Nutrients. 2015 Jan 8;7(1):390–404.

8. Kårlund A, Gómez-Gallego C, Korhonen J, Palo-oja OM, El-Nezami H, Kolehmainen M. Harnessing Microbes for Sustainable Development: Food Fermentation as a Tool for Improving the Nutritional Quality of Alternative Protein Sources. Nutrients. 2020 Apr 8;12(4):1020.

9. Guo Q, Chen P, Chen X. Bioactive peptides derived from fermented foods: Preparation and biological activities. Journal of Functional Foods. 2023 Feb;101:105422.

10. Peres Fabbri L, Cavallero A, Vidotto F, Gabriele M. Bioactive Peptides from Fermented Foods: Production Approaches, Sources, and Potential Health Benefits. Foods. 2024 Oct 23;13(21):3369.

11. Stipanuk MH, Caudill MA, editors. Biochemical, physiological, and molecular aspects of human nutrition. Fourth edition. St. Louis, Missouri: Elsevier; 2019. 959 p.

12. Chen M, Sun Q, Giovannucci E, Mozaffarian D, Manson JE, Willett WC, et al. Dairy consumption and risk of type 2 diabetes: 3 cohorts of US adults and an updated meta-analysis. BMC Med. 2014 Dec;12(1):215.

13. Soedamah-Muthu SS, Masset G, Verberne L, Geleijnse JM, Brunner EJ. Consumption of dairy products and associations with incident diabetes, CHD and mortality in the Whitehall II study. Br J Nutr. 2013 Feb 28;109(4):718–26.

14. Tapsell LC. Fermented dairy food and CVD risk. Br J Nutr. 2015 Apr;113(S2):S131–5.

15. Nikoloudaki O, Aheto F, Di Cagno R, Gobbetti M. Synthetic microbial communities: A gateway to understanding resistance, resilience, and functionality in spontaneously fermented food microbiomes. Food Research International. 2024 Sept;192:114780.

16. Kleiner M. Metaproteomics: Much More than Measuring Gene Expression in Microbial Communities. mSystems. 2019 June 25;4(3):e00115–19.

17. Van Den Bossche T, Armengaud J, Benndorf D, Blakeley-Ruiz JA, Brauer M, Cheng K, et al. The microbiologist’s guide to metaproteomics. iMeta. 2025 June;4(3):e70031.

18. Aydoğan C. Critical review of new advances in food and plant proteomics analyses by nano-LC/MS towards advanced foodomics. TrAC Trends in Analytical Chemistry. 2024 July;176:117759.

19. Hui YH, editor. Handbook of food and beverage fermentation technology. New York: Marcel Dekker; 2004. page 148. (Food science and technology).

20. Hui YH, Evranuz EÖ, editors. Handbook of Animal-Based Fermented Food and Beverage Technology. 2nd ed. Boca Raton: CRC Press; 2012.

21. Andrews S. FastQC: A Quality Control Tool for High Throughput Sequence Data

22. Bushnell B. BBTools software packag. e. 2014;

23. Gruber-Vodicka HR, Seah BKB, Pruesse E. phyloFlash: Rapid Small-Subunit rRNA Profiling and Targeted Assembly from Metagenomes Arumugam M, editor. mSystems. 2020 Oct 27;5(5):e00920–20.

24. Zougman A, Selby PJ, Banks RE. Suspension trapping (STrap) sample preparation method for bottom-up proteomics analysis. Proteomics. 2014 May;14(9):1006–1000.

25. Awan A, Bartlett A, Blakeley-Ruiz JA, Richie T, Theriot CM, Kleiner M. Dietary protein from different sources escapes host digestion and is differentially modified by the microbiota [Internet]. 2024 [cited 2024 July 29]. Available from: http://biorxiv.org/lookup/doi/10.1101/2024.06.26.600830

26. Mordant A, Kleiner M. Evaluation of Sample Preservation and Storage Methods for Metaproteomics Analysis of Intestinal Microbiomes Khursigara CM, editor. Microbiol Spectr. 2021 Dec 22;9(3):e01877–21.

27. Blakeley-Ruiz JA, Kleiner M. Considerations for constructing a protein sequence database for metaproteomics. Computational and Structural Biotechnology Journal. 2022;20:937–52.

28. Li W, Godzik A. Cd-hit: a fast program for clustering and comparing large sets of protein or nucleotide sequences. Bioinformatics. 2006 July 1;22(13):1658–9.

29. Bioinformatics solutions Inc. Bioinformatics solutions Inc.; 2019.

30. Vitorino R, Guedes S, Trindade F, Correia I, Moura G, Carvalho P, et al. *De novo* sequencing of proteins by mass spectrometry. Expert Review of Proteomics. 2020 Aug 2;17(7–8):595–607.

31. Zhang J, Xin L, Shan B, Chen W, Xie M, Yuen D, et al. PEAKS DB: De Novo Sequencing Assisted Database Search for Sensitive and Accurate Peptide Identification Molecular & Cellular Proteomics. 2012 Apr;11(4):M111.010587.

32. Dillies MA, Rau A, Aubert J, Hennequet-Antier C, Jeanmougin M, Servant N, et al. A comprehensive evaluation of normalization methods for Illumina high-throughput RNA sequencing data analysis. Briefings in Bioinformatics. 2013 Nov 1;14(6):671–83.

33. Kleiner M, Thorson E, Sharp CE, Dong X, Liu D, Li C, et al. Assessing species biomass contributions in microbial communities via metaproteomics. Nat Commun. 2017 Nov 16;8(1):1558.

34. Wickham H. ggplot2: elegant graphics for data analysis. Second edition. Cham: Springer international publishing; 2016. 1 p. (Use R!).

35. R Core Team. R: A language and environment for statistical computing. [Internet]. R Foundation for Statistical Computing, Vienna, Austria, 2023; Available from: https://www.R-project.org/.

36. Kolde R. Pheatmap: pretty heatmaps. R package version. 2019;1(2):726.

37. Mukherjee A, Breselge S, Dimidi E, Marco ML, Cotter PD. Fermented foods and gastrointestinal health: underlying mechanisms. Nat Rev Gastroenterol Hepatol. 2024 Apr;21(4):248–66.

38. Valentino V, Magliulo R, Farsi D, Cotter PD, O’Sullivan O, Ercolini D, et al. Fermented foods, their microbiome and its potential in boosting human health. Microbial Biotechnology. 2024 Feb;17(2):e14428.

39. Park I, Mannaa M. Fermented Foods as Functional Systems: Microbial Communities and Metabolites Influencing Gut Health and Systemic Outcomes. Foods. 2025 June 28;14(13):2292.

40. Aydoğan C. Critical review of new advances in food and plant proteomics analyses by nano-LC/MS towards advanced foodomics. TrAC Trends in Analytical Chemistry. 2024 May;117759.

41. Okeke ES, Ita RE, Egong EJ, Udofia LE, Mgbechidinma CL, Akan OD. Metaproteomics insights into fermented fish and vegetable products and associated microbes. Food Chemistry: Molecular Sciences. 2021 Dec;3:100045.

42. Yang L, Fan W, Xu Y. Metaproteomics insights into traditional fermented foods and beverages. Comp Rev Food Sci Food Safe. 2020 Sept;19(5):2506–29.

43. Tachie CYE, Onuh JO, Aryee ANA. Nutritional and potential health benefits of fermented food proteins. J Sci Food Agric. 2024 Feb;104(3):1223–33.

44. Tamang JP, Shin DH, Jung SJ, Chae SW. Functional Properties of Microorganisms in Fermented Foods Front Microbiol [Internet]. 2016 Apr 26 [cited 2025 May 7];7. Available from: http://journal.frontiersin.org/Article/10.3389/fmicb.2016.00578/abstract

45. Gavande PV, Goyal A, Fontes CMGA. Carbohydrates and Carbohydrate-Active enZymes (CAZyme): An overview. In: Glycoside Hydrolases [Internet]. Elsevier; 2023 [cited 2025 May 12]. p. 1–23. Available from: https://linkinghub.elsevier.com/retrieve/pii/B9780323918053000125

46. Di Rienzi SC, Britton RA. Adaptation of the Gut Microbiota to Modern Dietary Sugars and Sweeteners. Advances in Nutrition. 2020 May;11(3):616–29.

47. Kaoutari AE, Armougom F, Gordon JI, Raoult D, Henrissat B. The abundance and variety of carbohydrate-active enzymes in the human gut microbiota. Nat Rev Microbiol. 2013 July;11(7):497–504.

48. Lozupone CA, Stombaugh JI, Gordon JI, Jansson JK, Knight R. Diversity, stability and resilience of the human gut microbiota. Nature. 2012 Sept;489(7415):220–30.

49. Khan J, Zain WNIWM, Islam MN. Involvement of ACE2 in the intestinal transport of amino acids: Possible health and nutritional consequences in altered expression. Bangladesh J Med Sci. 2023 Sept 7;22(4):729–33.

50. Mirzaei M, Mirdamadi S, Ehsani MR, Aminlari M. Production of antioxidant and ACE-inhibitory peptides from Kluyveromyces marxianus protein hydrolysates: Purification and molecular docking. J Food Drug Anal. 2018 Apr;26(2):696–705.

51. Zhu XL, Watanabe K, Shiraishi K, Ueki T, Noda Y, Matsui T, et al. Identification of ACE-inhibitory peptides in salt-free soy sauce that are transportable across caco-2 cell monolayers. Peptides. 2008 Mar;29(3):338–44.

52. El Mecherfi KE, Todorov SD, Cavalcanti De Albuquerque MA, Denery-Papini S, Lupi R, Haertlé T, et al. Allergenicity of Fermented Foods: Emphasis on Seeds Protein-Based Products. Foods. 2020 June 16;9(6):792.

53. Günal-Köroğlu D, Karabulut G, Ozkan G, Yılmaz H, Gültekin-Subaşı B, Capanoglu E. Allergenicity of Alternative Proteins: Reduction Mechanisms and Processing Strategies. J Agric Food Chem. 2025 Apr 2;73(13):7522–46.

54. Huang X, Schuppan D, Rojas Tovar LE, Zevallos VF, Loponen J, Gänzle M. Sourdough Fermentation Degrades Wheat Alpha-Amylase/Trypsin Inhibitor (ATI) and Reduces Pro-Inflammatory Activity. Foods. 2020 July 16;9(7):943.

55. Landis EA, Oliverio AM, McKenney EA, Nichols LM, Kfoury N, Biango-Daniels M, et al. The diversity and function of sourdough starter microbiomes. eLife. 2021 Jan 26;10:e61644.

56. Reese AT, Madden AA, Joossens M, Lacaze G, Dunn RR. Influences of Ingredients and Bakers on the Bacteria and Fungi in Sourdough Starters and Bread Suen G, editor. mSphere. 2020 Feb 26;5(1):e00950–19.

57. De Vuyst L, Comasio A, Kerrebroeck SV. Sourdough production: fermentation strategies, microbial ecology, and use of non-flour ingredients. Critical Reviews in Food Science and Nutrition. 2023 June 11;63(15):2447–79.

58. Huys G, Daniel HM, De Vuyst L. Taxonomy and Biodiversity of Sourdough Yeasts and Lactic Acid Bacteria. In: Gobbetti M, Gänzle M, editors. Handbook on Sourdough Biotechnology [Internet]. New York, NY: Springer US; 2013 [cited 2025 Nov 6]. p. 105–54. Available from: https://link.springer.com/10.1007/978-1-4614-5425-0_5

59. Urien C, Legrand J, Montalent P, Casaregola S, Sicard D. Fungal Species Diversity in French Bread Sourdoughs Made of Organic Wheat Flour. Front Microbiol. 2019 Feb 18;10:201.

60. García-Béjar B, Fernández-Pacheco P, Carreño-Domínguez J, Briones A, Arévalo-Villena M. Identification and biotechnological characterisation of yeast microbiota involved in spontaneous fermented wholegrain sourdoughs. J Sci Food Agric. 2023 Dec;103(15):7683–93.

61. Perez-Riverol Y, Bai J, Bandla C, García-Seisdedos D, Hewapathirana S, Kamatchinathan S, et al. The PRIDE database resources in 2022: a hub for mass spectrometry-based proteomics evidences. Nucleic Acids Research. 2022 Jan 7;50(D1):D543–52.

